# Nonstructural protein 1 of swine arterivirus PRRSV downregulates promyelocytic leukemia nuclear bodies and promotes viral replication

**DOI:** 10.1101/2023.08.04.552021

**Authors:** Chia-Ming Su, Mingyuan Han, Roger Everett, Dongwan Yoo

## Abstract

Porcine reproductive and respiratory syndrome virus (PRRSV) inhibits the type I interferon (IFN) production and signaling pathways during infection, and the nonstructural protein 1 (nsp1) has been identified as a potent IFN antagonist. Promyelocytic leukemia (PML) protein is a major scaffold protein organizing the PML nuclear bodies (NBs) of the cell and plays a diverse role in maintaining the cellular homeostasis including antiviral response among many other processes. The present study reveals a significant reduction of PML NBs in cells during infection of PRRSV, implicating the negative regulation of PML gene expression by PRRSV. Subsequently, the nsp1β protein was identified as the viral regulator for PML expression. The overexpression of PML isoforms restricted viral replication, while the gene silencing of endogenous PML promoted viral replication. The downregulation of PML expression by PRRSV was post-translational via the ubiquitination-proteasome pathway. Of six isoforms, PML-II and PML-IV exhibited the most potent suppressive activity against viral replication. PRRSV nsp1β bound to PML directly, and this interaction was mediated through the small ubiquitin-like modifier (SUMO)-interacting motifs (SIMs) on nsp1β. Further studies revealed that double mutations in SIM1 and SIM4 abolished the binding of nsp1β to PML and prevented the PML degradation. The PML downregulation by nsp1 was common in other arteriviruses including equine arteritis virus, murine lactate dehydrogenase elevating virus, and simian hemorrhagic fever virus. Our study unveils the evolutionary conservation of the viral immune evasion strategy employed by arteriviruses, which promotes their replication by targeting PML for downregulation.

**IMPORTANCE OF THE STUDY:** Porcine reproductive and respiratory syndrome (PRRS) is an economically significant disease in the swine industry worldwide. One of the immunological hallmarks in virus-infected animals is the suppression of type I interferon response during an early-stage infection, leading to the consequence of adaptive immunity and viral persistence. In the present study, we report that the nsp1-beta protein of PRRS virus degrades the promyelocytic leukemia (PML) protein and downregulates PML nuclear body (NB) formation. The PML downregulation by PRRS virus results in enhanced viral replication. The PML downregulation by nsp1 is common in other arteriviruses, unveiling the basic understanding of cell-virus interactions and immune evasion strategies for arteriviruses.

## INTRODUCTION

Porcine reproductive and respiratory syndrome virus (PRRSV) is a single-stranded positive-sense RNA virus belonging to the family *Arteriviridae* in the order *Nidovirales*. The *Arteriviridae* family consists of 6 subfamilies (*Equarterivirinae*, *Simarterivirinae*, *Variarterivirinae*, *Zealarterivirinae*, *Heroarterivirinae*, and *Crocarterivirinae),* and within the family, there are 13 genera and 23 species according to the most recent version of the International Code of Virus Classification and Nomenclature (https://talk.ictvonline.org/taxonomy). The most studied and frequently sampled arteriviruses are represented by PRRSV, equine arteritis virus (EAV), lactate dehydrogenase elevating virus (LDV) of mice, and simian hemorrhagic fever virus (SHFV). PRRSVs are categorized into two species, *Betaarterivirus suid* 1 (PRRSV-1) and *Betaarterivirus suid* 2 (PRRSV-2), based on their genomic sequence similarities. The PRRSV genome varies from 14.9 to 15.5 kb in length and consists of the 5’ untranslated region (UTR), 11 open reading frames (ORFs) followed by the 3’ UTR and a polyadenylated tail (1). ORF1a and ORF1b code for two large polyproteins but share a single translational start so that ORF1b is expressed as a fusion protein with ORF1a by ribosomal frame shafting. The two polyproteins are further processed to at least 16 nonstructural proteins (nsps) by four viral proteinases (2). The remaining ORFs produce eight structural proteins: E, GP2, GP3, GP4, GP5, ORF5a, M, and N.

Type I interferons (IFNs-α/β) play a pivotal role as antiviral cytokines in the initial phase of viral infection. When cells are infected with a virus, the IFN signaling is triggered, and a series of IFN-stimulated genes (ISGs) are expressed, contributing to the establishment of an antiviral state of the cells (3). For PRRSV infection, however, a notable characteristic has been observed for the poor production of type I IFNs, and eight viral proteins (nsp1α, nsp1β, nsp2, nsp2TF, nsp2N, nsp4, nsp11, and N) have been identified to participate in the blocking of IFN production and signaling (4–13). Among these proteins, nsp1, comprising the nsp1α and nsp1β subunits, exhibit the most potent IFN antagonistic property. The nsp1α subunit inhibits the production of type I IFNs and impedes the promoter activity when stimulated with the dsRNA analogue (14). It also downregulates IFN production by degrading the CREB (cyclic-AMP-responsive element binding)-binding protein (CBP) through the proteasomal pathway (15). Furthermore, PRRSV nsp1α suppresses NF-κB activation in a RIG-I-dependent manner, leading to the suppression of type I IFN production (12, 16). The nsp1β subunit has also been identified as an effective IFN antagonist (17). PRRSV nsp1β inhibits the STAT1 (signal transducers and activators of transcription 1) phosphorylation and disrupts the nuclear translocation of ISGF3 (interferon stimulated gene factor 3), thereby suppressing the JAK (Janus kinase)-STAT signaling pathway and inhibiting the expression of ISGs (10, 14). In addition, nsp1β degrades karyopherin-α1 (KPNA1), a nuclear transporter protein required for ISGF3 nuclear translocation (18). A recent study shows that nsp1β binds to nucleoporin 62 (Nup62) and disrupts the nuclear pore complex (NPC). The disruption of NPC results in the blocking of nucleocytoplasmic export of host mRNAs and the reducing of host protein synthesis including type I IFNs, ISGs, and IRF3 (19). The SAP motif [SAF-A/B (scaffold attachment factors A and B), Acinus, and PIAS (protein inhibitor of activated STAT] has been identified in nsp1β and is the critical domain for host mRNA nuclear retention and IFN suppression (20). The combined activities of the nsp1α subunit primarily targeting the IFN production and the nsp1β subunit affecting both IFN production and signaling make the nsp1 protein the potent type I IFN antagonist of PRRSV.

Promyelocytic leukemia nuclear bodies (PML-NBs) are dynamic cellular structures that play a key role in diverse cellular processes including stress response, gene regulation, oncogenesis (21), cell senescence, DNA damage repair, apoptosis, and antiviral defense (for reviews, see Everett et al., 2013; Everett and Chelbi-Alix, 2007; Neerukonda, 2021). The size, number, and localization of PML-NBs can vary depending on the cell type, cell cycle phase, and physiological state. The main scaffold protein of PML-NBs is the promyelocytic leukemia (PML) protein now known as the tripartite motif 19 (TRIM19). TRIM is a family of proteins characterized by the presence of a highly conserved RBCC motif made of tandemly arranged RING finger domain (R), cysteine-histidine-rich B box domains (B1 and B2), and an alpha-helical coiled-coil domain (CC) (25, 26). Alternative mRNA splicing gives rise to seven different isoforms of PML (I through VII) which possess the identical RBCC motifs in their N-termini but differ in their C-termini (27, 28). PML-VII lacks the C-terminal nuclear localization signal and thus remains cytoplasmic (27). PML is subject to SUMOylation which involves the covalent attachment of small ubiquitin-like modifier (SUMO) to lysine (K) at specific sites. PML also contains SUMO interacting motifs (SIMs) that allows them to interact with SUMOylated client proteins (29). PML as the scaffold organizer of PML-NBs functions through SUMO-SIM interactions. Other components of PML-NBs include Sp100, DAXX (death-domain associated protein), and ATRX, which acts as intrinsic restriction factors that repress viral replication (30–32).

PML is an interferon-stimulated gene (ISG) product. When cells are infected with a virus, the number and size of PML-NBs increase in response to IFN signaling. PML associates with transcription factors NF-κB, STAT1, and CBP, and facilitates the formation of transcriptional complexes on the promoters of IFN-β and multiple ISGs (33). PML can also potentiate IRF3 induction by sequestering PIN1 [peptidylprolyl cis/trans isomerase, NIMA (never in mitosis A)-interacting 1] to PML-NBs, thus preventing the PIN1-mediated downregulation of IRF3 via ubiquitin-dependent degradation (34, 35). PML inhibits not only DNA viruses but also RNA viruses. PML knock-out mice become susceptible to infections by rabies virus and lymphocytic choriomeningitis virus (36, 37). Consequently, viruses seem to have developed a particular strategy to evade the antiviral activity of PML.

In the present study, we showed that PRRSV downregulated the PML expression to evade host cell antiviral defense. The PRRSV nsp1β protein bound directly to PML and mediated PML degradation through the ubiquitin-proteasome pathway. The PML degradation promoted efficient viral replication, which is an important strategy of arteriviruses for immune evasion and enhanced viral replication.

## RESULTS

### Reduction of PML-NBs in PRRSV-infected cells

The PML protein is one of ISGs and has emerged as a significant regulator for viral replication. Conversely, many viruses have evolved to counteract the antiviral effects of PML. To determine whether PRRSV possessed the ability to counteract the PML function during infection, MARC-145 cells were infected with PRRSV for 24 h and stimulated with IFNs, given that PML is an IFN-inducible gene. The potential of PRRSV for PML regulation was then examined by staining with α-PML antibody and α-PRRSV N antibody. After stimulation, numbers of PML-NBs in the nucleus increased as expected (Fig. 1A), while their numbers in un-stimulated cells remained minimal. In contrast, numbers of PML-NBs in virus-infected cells decreased significantly even after IFN stimulation (long white arrows) compared to those in un-infected cells (short yellow arrows). The decrease of PML-NBs was quantified by counting the numbers of PML-NB puncta per cell for 20 cells (Fig. 1B). A considerable decrease in the number of PML-NBs was observed in virus-infected cells compared to un-infected cells. The decrease was pronounced with the average number of 3.45 PML-NBs/cell compared to the average number of 12.22 PML-NBs/cell, suggesting a possible downregulation of PML-NBs by PRRSV. To confirm the PRRSV-mediated PML downregulation, the levels of PML expression were determined at various times post-infection (hpi) by Western blot. The relative expression of PML protein was represented by the density of band with respect to that of the actin-normalized control, and the mock-infection group was set as 1 (Fig. 1C). The PRRSV infection led to the reduced expression of PML to 0.78 at 24 h hpi, and the PML remained decreased throughout the infection. Our results demonstrate that PRRSV downregulated PML expression and disrupted the PML-NB formation in the nucleus, suggesting a potent of PRRSV for IFN antagonism.

**Fig. 1.**
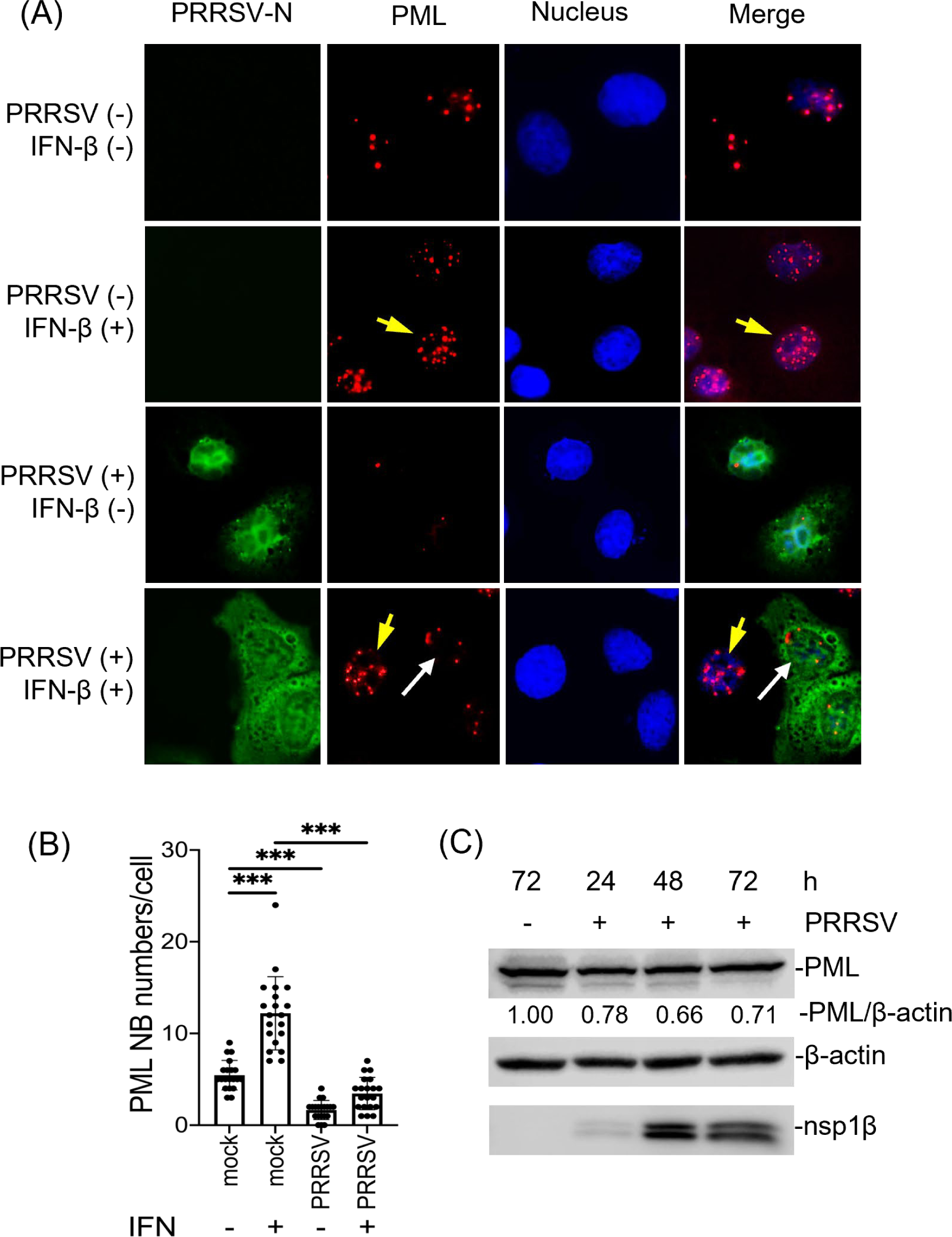
Reduction of PML-NBs in PRRSV-infected cells. (A), MARC-145 cells were infected with the PA8 strain of PRRSV at 1 moi for 24 h and incubated with IFN-β (1,000 unit/ml) for 6 h. Cells were stained with α-PRRSV-N MAb (MR40) (mouse) (green) and α-PML PAb (rabbit) (red). Nuclei were stained with DAPI (blue). Long white arrows indicate infected cell, and short yellow arrows indicate uninfected cell. Images were taken by confocal microscopy (Nikon A1R). (B), PML-NB numbers per cell. The PML-NB puncta were counted from randomly chosen 20 cells. Statistical significance is indicated as follows: ***P<0.01. (C), MARC-145 cells were infected with the PA8 strain of PRRSV at 1 moi for 24 h and cell lysates were subjected to Western blot using α-PML PAb. β-actin was used as a loading control. Intensities of PML staining were quantified with the Image J system, normalized to that of β-actin, and compared to the mock control. Numbers each band indicate relative fold changes.

### Downregulation of PML expression by PRRSV nsp1β protein

The nsp1 protein of PRRSV is the most potent IFN antagonist. The nsp1β protein downregulates the nucleocytoplasmic trafficking of host mRNAs and inhibits the type I IFNs and ISG transcriptions (19, 38). Since PML functions as an ISG, it was of interest to examine if the PRRSV nsp1 protein caused the reduction of PML in virus-infected cells. We thus first examined the PML-NB formation in nsp1-expressing cells by IFA. The ORF3a protein of SARS-CoV-2 was previously shown to be irrelevant to IFN response (39) and so was included as a negative control. After stimulation with IFN, the numbers of PML-NBs were decreased in nsp1α-expressing and nsp1β-expressing cells (long white arrows) compared to control (short yellow arrows) (Fig. 2A). The decrease in the number of PML-NBs was evident in both nsp1α-expressing and nsp1β-expressing cells even after IFN stimulation (Fig. 2B). The PML mRNA was assessed by RT-qPCR in nsp1 gene-transfected cells. The amount of mRNA, however, was not altered in nsp1α-expressing nor nsp1β-expressing cells (Fig. 2C), suggesting that the PML downregulation could be a post-translational event without impeding transcription. Thus, the PML protein was determined by Western blot (Fig. 2D, 2E). Even after stimulation, the reduction of PML was significant to the levels of 0.84 and 0.78 in cells expressing nsp1α and nsp1β, respectively, in comparison to that of control cells (Fig. 2E). Our findings indicate that the PML downregulation is post-translational and that nsp1α and nsp1β are the viral components reducing the PML expression.

**Fig. 2.**
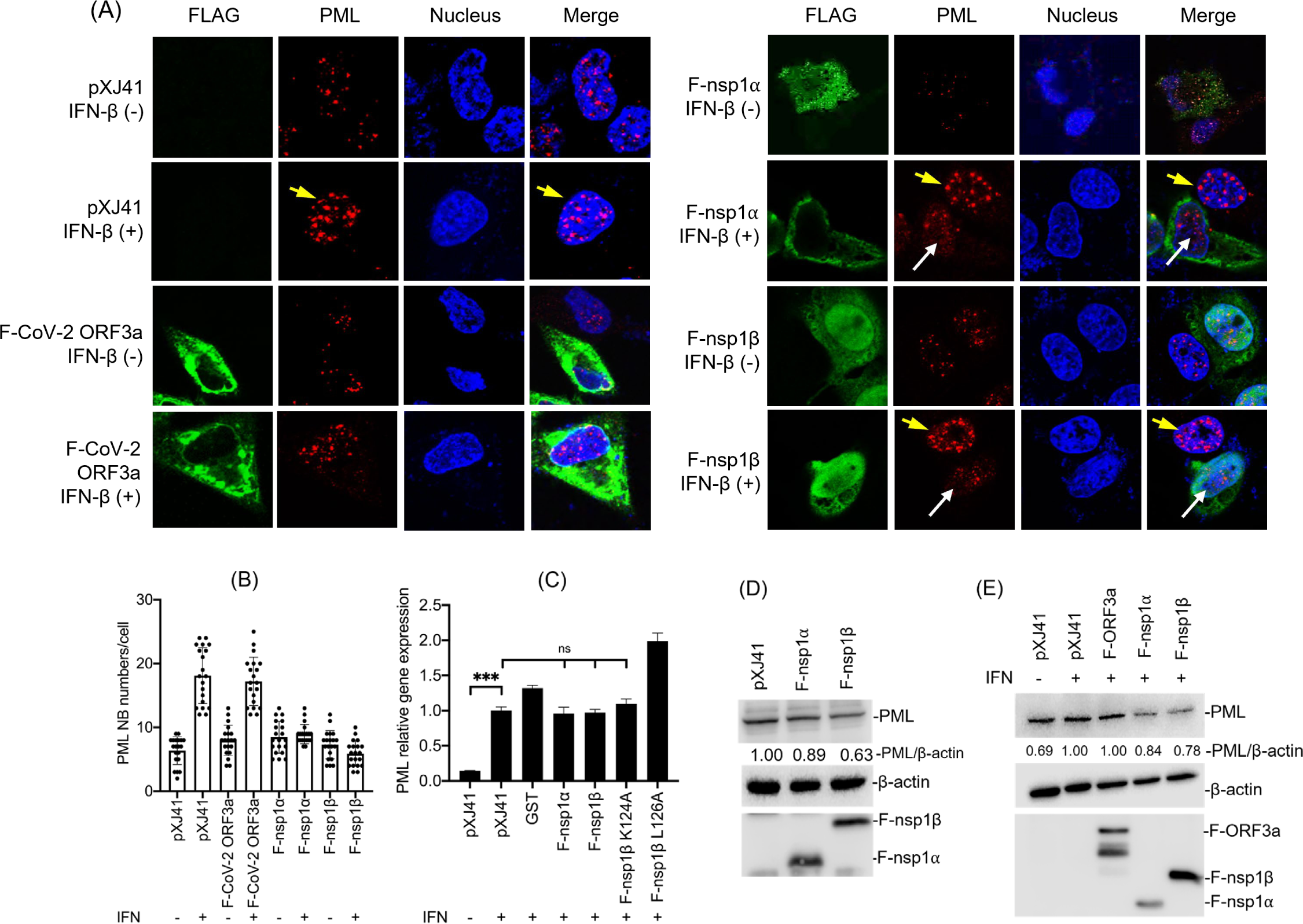
PRRSV nsp1-mediated PML downregulation. (A), Reduction of PML-NB numbers by PRRSV nsp1. HeLa cells were transfected with indicated genes and incubated with IFN-β (1000 unit/ml). The cells were stained with α-FLAG MAb (green) and α-PML PAb (red). Nuclei were stained with DAPI (blue). Long white arrows indicate gene transfected cells, and short yellow arrows indicate untransfected cell. The images were taken by confocal microscopy (Nikon A1R). (B), RT-qPCR for PML protein genes in nsp1-expressing cells. HeLa cells were transfected with 2 μg of indicated genes for 24 h and incubated with IFN-β (1,000 unit/ml) for 6 h followed by RT-qPCR using the total cellular RNA. The expression levels were calculated with 2^−ΔΔCT^ method by normalizing to that of β-actin. The fold changes were calculated with respect to the level of pXJ41. Error bars, mean ± standard deviation (s.d.). ns: no significant difference. ***, P<0.001. (C), The PML-NB numbers per nucleus for 20 cells. (D, E) HeLa cell lysates expressing indicated gene were subjected to Western blot using α-PML PAb. β-actin was used as a loading control. Intensities of PML staining were quantified with the ImageJ system (NIH), normalized to that of β-actin, and compared to the mock control. Numbers around each band indicate relative fold changes.

### PML degradation was independent from PLP1β and SAP motifs of nsp1β

Since PRRSV nsp1β contains the papain-like proteinase (PLP) 1β domain, the PLP1β was postulated to cause PML degradation. For encephalomyocarditis virus, 3C^pro^ alters the cellular localization of PML and reduces its expression (40). Similarly, 3C^pro^ of enterovirus 71 cleaves isoforms of PML and results in their reduction (41). For PRRSV, the PLP1β domain contains a SAP motif (Fig. 3A), and leucine 126 (L126) in SAP has been shown to be essential for IFN antagonism (20, 38). This information led us to examine whether those two functions of nsp1β were associated with PML downregulation. To examine this, we generated four mutants; two for the SAP motif and two for the PLP1β motif. L126A represented the leucine-to-alanine mutant at position 126 to knock-out the SAP function, and K124A was the lysine-to-alanine mutant to serve as a SAP control since lysine 124 is nonessential for the SAP function. For PLP1β, C90 and H159 are catalytic residues for the proteinase activity, and thus, we generated the C90A and H159A mutants to knock-out the papain-like proteinase activity. We expressed the four mutant genes in cells and determined the numbers of PML-NBs by IFA. A significant reduction in numbers of PML-NBs was still observed in cells expressing each of the SAP and PLP1β mutants (Fig. 3B). The reductions were quantified by counting PML-NBs, and the results confirmed that the loss of the SAP and PLP1β functions did not restore the PML-NB formation (Fig. 3C). These findings demonstrated that the reduction of PML-NBs by nsp1β was independent from its PLP1β activity or SAP-mediated IFN antagonism. Intriguingly, however, PML was found to co-localize with nsp1β (Fig. 3D), suggesting a possible interaction of PML with nsp1β in the nucleus.

**Fig. 3.**
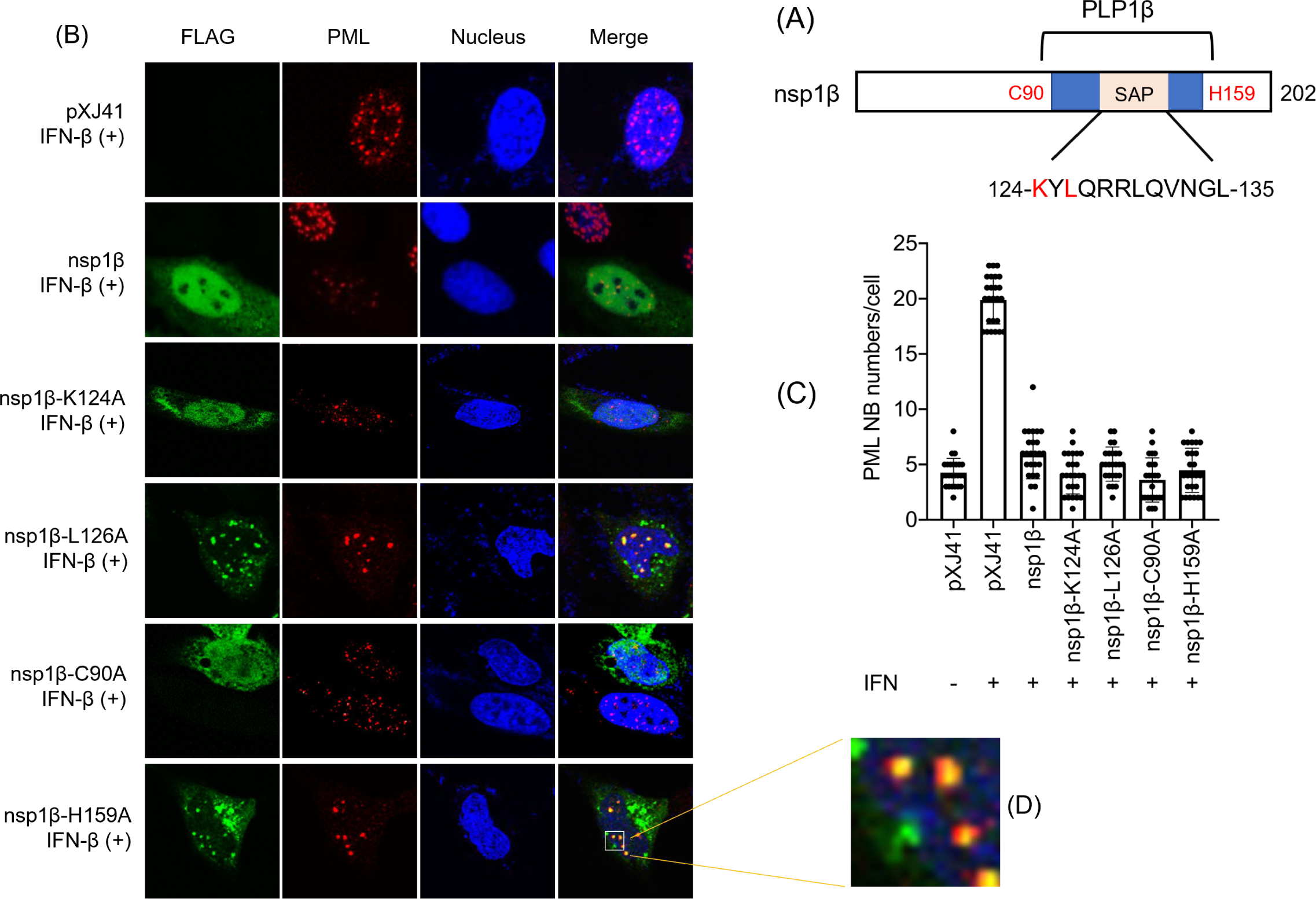
The PML degradation is independent from the PLP1β and SAP motifs of nsp1β. (A) Depiction of functional domains in PRRSV nsp1β. Blue box indicates papain-like proteinase (PLP1β) domain, and pink box indicates SAP motif. Red color indicates amino acid mutations. Numbers indicate amino acid positions. (B, D), HeLa cells were expressed with indicated nsp1β mutants and stimulated with IFN-β (1,000 unit/ml). The cells were stained with α-FLAG MAb (green) and α-PML PAb (red). Nuclei were stained with DAPI (blue). The images were taken by confocal microscopy (Nikon A1R). (C), The numbers of PML-NBs per nucleus counted from 20 cells.

### PML was degradated via ubiquitin-proteasome pathway

Previously, PRRSV nsp1 was shown to hinder type I IFN production by degrading the CREB-binding protein in the nucleus (15). This process did not require the nsp1 proteinase activity but implicated the involvement of ubiquitine-proteasomal degradation pathway. Since the PLP1β activity of nsp1β was not involved in the PML downregulation (Fig. 3), it was plausible that the ubiquitine-proteasomal pathway might trigger PML degradation. To examin this, MG132, a potent inhibitor of ubiquitin proteasome-dependent degradation was used. HeLa cells were transfected with the nsp1α or nsp1β gene, and at 24 h, treated with MG132 for 6 h followed by the staining for PML. After MG132 treatment, the number of PML-NBs was significantly increased in F-nsp1β-expressing cells (Figs. 4A, white arrow; 4B). Interestingly, F-nsp1α-expressing cells did not restore the PML numbers even with MG132 treatment (Fig. 4B). Western blot analysis confirmed that MG132 blocked the PML degradation in F-nsp1β-expressing cells but not in nsp1α-expressing cells (Fig. 4C). These findings indicate that the PML reduction was mediated by nsp1β via the ubiquitin-proteasome pathway and that nsp1α may function differently from nsp1β for PML degradation. We did not pursue the basis for nsp1α-mediated PML degradation further in the present study but examined the nsp1β-mediated ubiquitination of PML using the HA-Ubiquitin (Ubi) construct. Cells were co-expressed with F-nsp1β and HA-Ubiquitin, and cell lysates were incubated with α-PML Ab, followed by immunoblot using α-HA Ab. As shown in Fig. 4D, the ubiquitination of PML was strongly enhanced in HA-Ubi- and nsp1β-coexpressing cells (left panel). The ubiquitination of endogenous PML was also evaluated using α-Ubi Ab, and the data showed an increased level of ubiquitinated PML in F-nsp1β-expressing cells compared to control (Fig. 4E, left panel). Taken together, our data demonstrate that the PRRSV nsp1β protein mediates the PML degradation via ubiquitin-proteasome pathway.

**Fig. 4.**
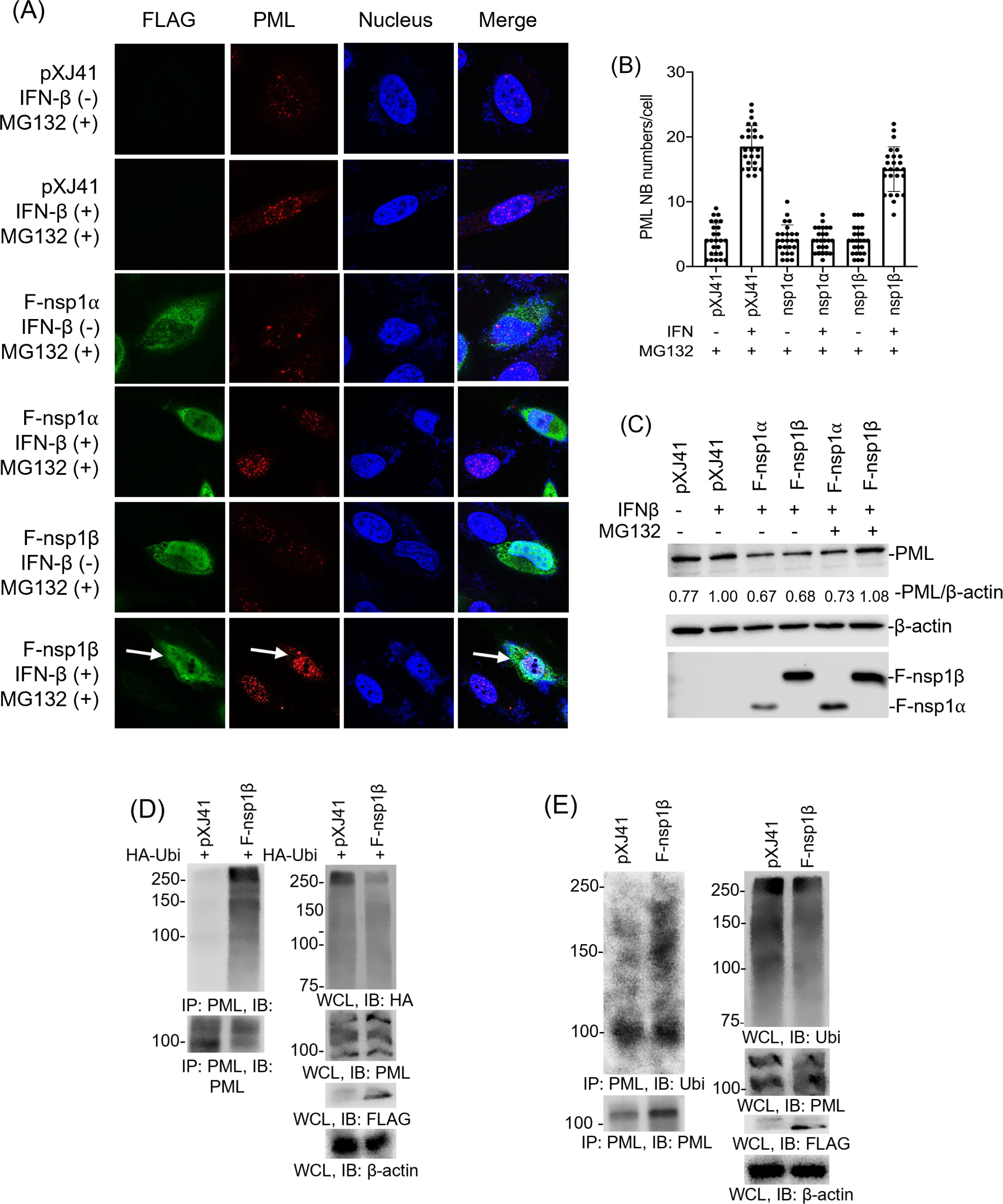
Degradation of PML protein by nsp1β through ubiquitin-proteasome pathway. (A), MG132 treatment restores the PML-NBs formation in nsp1β expressing cells. HeLa cells were expressed with indicated genes and treated with IFN-β (1000 unit/ml) and MG132 (10 μM). The cells were stained with α-FLAG MAb (green) and α-PML PAb (red). Nuclei were stained with DAPI (blue). White arrows indicate gene-transfected cells, and yellow arrow indicates un-transfected cell. The images were taken by confocal microscopy (Nikon A1R). (B), The PML-NB numbers per cell. The PML-NB puncta were counted from 20 cells. (C), HeLa cells were expressed with indicated genes and treated with IFN-β (1000 unit/ml) and MG132 (10 μM). Cell lysates were prepared and subjected to Western blot analyses using α-PML PAb. β-actin was used as a loading control. Intensities of PML staining were quantified with the Image J system, normalized to that of β-actin, and compared to the mock control. Numbers indicate relative fold changes. HeLa cells were transfected with FLAG-nsp1β and HA-Ubi (D), or FLAG-nsp1β alone (E) for 24 h, and cell lysates were prepared and subjected to co-IP using α-PML PAb for immunoprecipitation and α-PML PAb, α-HA MAb (D) or α-Ubi MAb (E) for immunoblot (left panels). Whole cell lysates were subjected to Western blot as an input control (right panels).

### Interaction of PRRSV nsp1β with PML

Since PML colocalized in puncta with nsp1β (Fig. 3D), the direct binding of two proteins was examined by GST (glutathione S-transferase) pulldown assays. PRRSV nsp1β was expressed as a GST fusion protein in *Escherichia coli*, and the resulting protein was immobilized to GST-coupled Sepharose beads. The beads were incubated with whole cell lysates, and bound proteins were detected by immunoblot using α-PML Ab. The GST-nsp1β fusion protein specifically pulled down the PML protein (Fig. 5A, upper panel), providing solid evidence for the binding of nsp1β to endogenous PML. To further confirm their finding, co-immunoprecipitation (co-IP) was conducted. F-nsp1β and PML were over-expressed in cells, and the lysates were incubated with α-PML Ab, followed by immunoblot with α-FLAG Ab (Fig. 5B). It was evident that F-nsp1β was co-immunoprecipitated with PML. To examine if specific isoforms of the PML protein interacted with nsp1β, six individual isoforms of HA-PML was co-expressed with F-nsp1β, and each lysate was co-immunoprecipitated using α-HA Ab and immunoblot using anti-FLAG Ab. All six isoforms of PML-I through PML-VI were interactive with F-nsp1β (Fig. 5C). Our data showed that, regardless of the endogenous or exogenous PML, nsp1β bound to all six isoforms of PML.

**Fig. 5.**
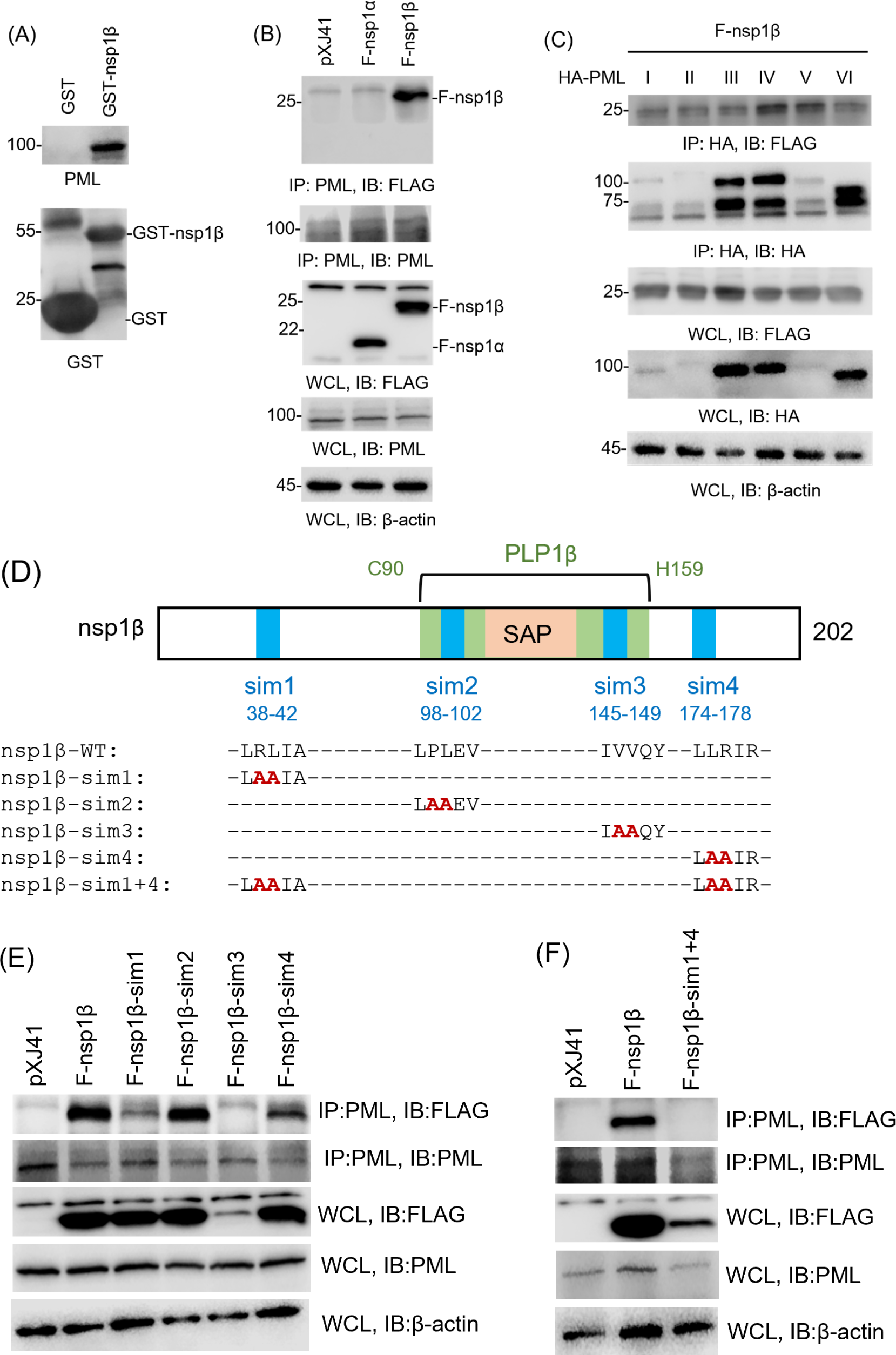
Interaction of nsp1β SIM motif with PML protein. (A), GST pulldown assay for PML. GST or GST-nsp1β was expressed in *E. coli* and coupled with glutathione-Sepharose 4B beads for incubation with HeLa cell lysates. The precipitates were subjected to Western blot using α-PML PAb. (B, C) Specific binding of nsp1β with endogenous PML (B), and six different isoforms of PML (PMLI - IV) expressed by gene transfection. Their bindings were determined by co-IP. HeLa cells were co-transfected with 2 μg of indicated genes and each of HA-PMLs I through IV (2 μg) for 24 h and were subjected to co-IP. Cells lysates were pulled down by α-PML PAb (B) or α-HA MAb (C) and probed with α-FLAG PAb. Whole cell lysates (WCL) were subjected to Western blot as an input control. (D), Schematic representation of nsp1β SUMO interacting motif and introduction of mutations. Potential SIM motifs were predicted using the online tool for GPS-SUMO prediction (Zhao et al., 2014; https://sumo.biocuckoo.cn/). (E, F), HeLa cells were transfected with 2 μg of indicated plasmid and incubated for 24 h. Cell lysates were prepared and subjected to co-IP with α-PML PAb, followed by Western blot using α-FLAG PAb. WCL were probed with α-PML PAb or α-FLAG PAb. β-actin served as a loading control.

The physical stability of PML-NBs relies on the SUMOylation of PML. SUMOylated PML recruits client proteins containing a SUMO-interacting motif (SIM) to the NB inner core via SUMO-SIM interactions (25). Since PRRSV nsp1β bound to all six isoforms of PML (Fig. 5C), we hypothesized that the nsp1β binding to PML would possibly be mediated through SUMO-SIM interactions. We first examined the nsp1β sequence by bioinformatics analysis (42), and four high-confidence SIM motifs were predicted with the sequences of _38_-LRLIA-_42_ (named sim1; numbers in subscript indicate positions), _98_-LPLEV-_102_ (sim2), _145_-IVVQY-_149_ (sim3), and _174_-LLRIR-_178_ (sim4) (Fig. 5D). The predicted motifs were then mutated by introducing two alanines to respective SIMs, followed by expression and co-IP to determin their interactions with PML. Of four SIM mutant constructs, nsp1β-sim1 and nsp1β-sim4 resulted in the decreased binding to PML, compared to wild-type nsp1β and nsp1β-sim2 (Fig. 5E). Thus, both SIM1 and SIM4 of nsp1β were postulated to interact with SUMO on the PML protein. F-nsp1β-sim3 also abolished its binding to PML (Fig. 5E). However, the ectopic expression of nsp1β-sim3 was notably low in the whole cell lysate compared to that of wild-type nsp1β. Thus, the loss of its binding to PML might be attributed to the low level expression of nsp1β-sim3 and its peculia cellular distribution (data not shown), and so this construct was disregarded in the further study. To confirm SIM1 and SIM4 as the PML binding motifs, dual mutations were introduced to generate F-nsp1β-sim1+4, followed by co-IP. The F-nsp1β-sim1+4 dual mutant resulted in the complete loss of binding to PML (Fig. 5F), demonstrating that both SIM1 and SIM4 are crucial motifs for the binding of nsp1β to PML.

### PML degradation by nsp1β resulted in IFN downregulation

SIM plays a vital role in the regulation of PML expression and viral pathogenesis (43, 44). Since SIM1 and SIM4 of nsp1β was determined as crucial motifs for PML binding, it was of interest to examine the virological role of the nsp1β-binding and PML degradation. We first examined the PML-NB formation in SIM mutant nsp1β-expressing cells. While the number of PML-NBs decreased in wild-type nsp1β-expressing cells, the numbers of PML-NBs were increased in the nsp1β-sim1-expressing and nsp1β-sim1+4-expressing cells (Fig. 6A). The increased expressions of PML in SIM mutant-expressing cells were further confirmed by Western blot. As anticipated, PML expressions were elevated in the nsp1β-sim1- and nsp1β-sim3-expressing cells (Fig. 6B). Furthermore, cells expressing nsp1β-sim1+4 restored the PML expression to the level of control, whereas PML expression was decreased in cells expressing wild-type nsp1β (Fig. 6C). These results demonstrate that the interaction between PML and nsp1β causes the degradation of PML protein.

**Fig. 6.**
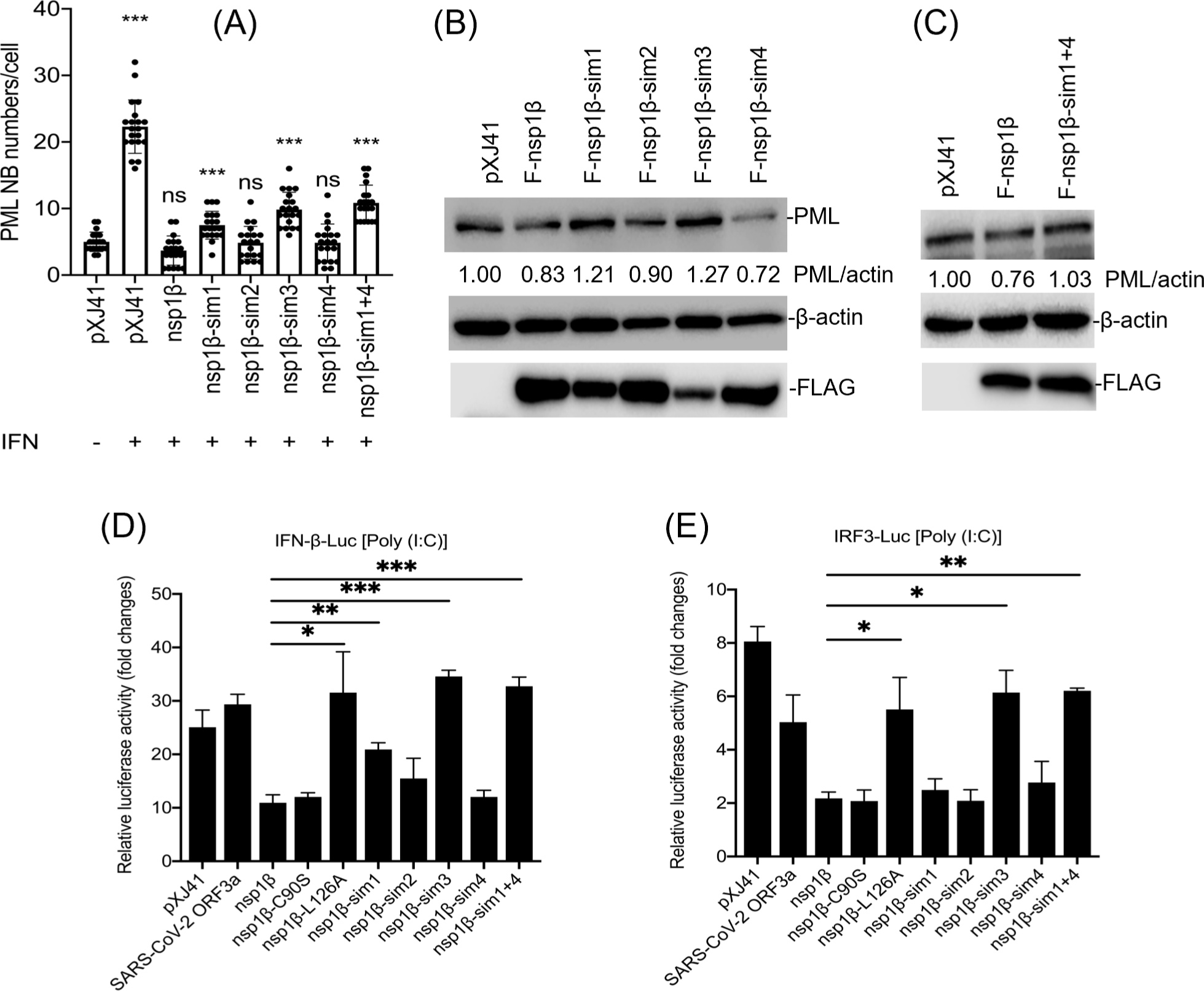
The interaction between PML and nsp1β results in PML downregulation. (A), Numbers of PML-NBs in indicated gene-expressing cells. PML-NBs formation was restored in SIM mutant nsp1β-expressing cells. HeLa cells were expressed with indicated genes and stimulated with IFN-β (1000 unit/ml), followed by staining with α-FLAG MAb (green) and α-PML PAb (red). The PML-NB puncta were counted from 20 cells. (B, C), Levels of PML protein in SIM mutant nsp1β-expressing cells. HeLa cells-expressing indicated gene were subjected to Western blot using α-PML PAb. β-actin was used as a loading control. Intensities of PML staining were quantified with the Image J system, normalized to that of β-actin, and compared to the mock control. Numbers indicate relative fold changes. (D, E), Regulation of type I IFN production by SIM mutant nsp1β. HeLa cells were co-transfected with pRL-TK and pIFN-β-luc (D) or pIRF3-luc (E) along with respective viral gene and stimulation with poly(I:C) for 12 h. The firefly and Renilla luciferase activities were determined. Error bars, mean ± standard deviation (s.d.). *, P<0.05; **, P<0.01; ***, P<0.001.

PML plays a pivotal role in the activation of innate immunity against viral infections by facilitating the formation of transcription complex on the IFN-β gene promoter via specific association with NF-κB and STAT1 (33). Thus, it was of interest to examine the IFN response to nsp1β-PML interaction using the IFN-β reporter assays. Cells were co-expressed with each of the SIM mutant and pIFN-β-luc, and the reporter expressions were quantified. While the reporter expression was increased by poly(I:C) stimulation in control cells, it was significantly reduced in wild-type nsp1β-expressing cells as expected (Fig. 6D). The L126A SAP mutant demonstrated higher reporter expression compared to wild-type nsp1β after poly(I:C) stimulation. Notably, F-nsp1β-sim3 and F-nsp1β-sim1+4 elicited the reporter expression higher than wild-type nsp1β (Fig. 6D), indicating that the SIM mutations resulted in the loss of nsp1β-mediated IFN suppression. Previously, PML was shown to enhance the induction of IFN-β by IRF3 activation (35). Thus, the SIM mutants were examined for their ability of IRF3 suppression. The pIRF3-luc construct contains 4 copies of the IRF3 binding sequence in front of the luciferase reporter gene, and so the increase of reporter expression represents the IRF3 activation. While nsp1β exhibited a strong suppression of IRF3 response, F-nsp1β-sim3 and F-nsp1β-sim1+4 showed minor suppressions (Fig. 6E). Together, these results demonstrate that the PML degradation mediated by nsp1β is linked to the suppression of IRF3 activity, subsequently leading to the IFN downregulation.

### PML protein suppressed PRRSV infection

PML has been shown to possess an antiviral activity against a number of viruses (24, 45–48). To determine if PML was also antiviral against PRRSV, we examined the viral replication in PML knock-down cells by gene silencing. The PML-specific small interfering RNA (siPML) targeting all six isoforms efficiently downregulated PML mRNA in MARC-145 cells (Fig. 7A). Western blot verified 71% reduction of PML transcript by siPML gene-silencing compared to the control at 24 h after treatment (Fig. 7B). The role of PML on PRRSV replication was then assessed by determining viral replication by RT-qPCR for viral N-specific mRNA and TCID_50_ (tissue culture infectious dose 50) in PML gene-silencing cells. The N-specific mRNA became increased during 24 to 72 hpi compared to siControl (Fig. 7C). The viral titers were also higher in PML knock-down cells as early as 24 hpi (Fig. 7D). The antiviral activity of PML protein was further confirmed by over-expressing PML. PMLs I through VI were co-expressed in MARC-145 cells for 48 h which were then infected with PRRSV. In PML over-expressing cells, the viral N-specific mRNA was decreased, and the viral titers were also decreased compared to the control at 48 hpi and 72 hpi (Figs. 7F, 7G), demonstrating that PML possessed the antiviral activity against PRRSV replication.

**Fig. 7.**
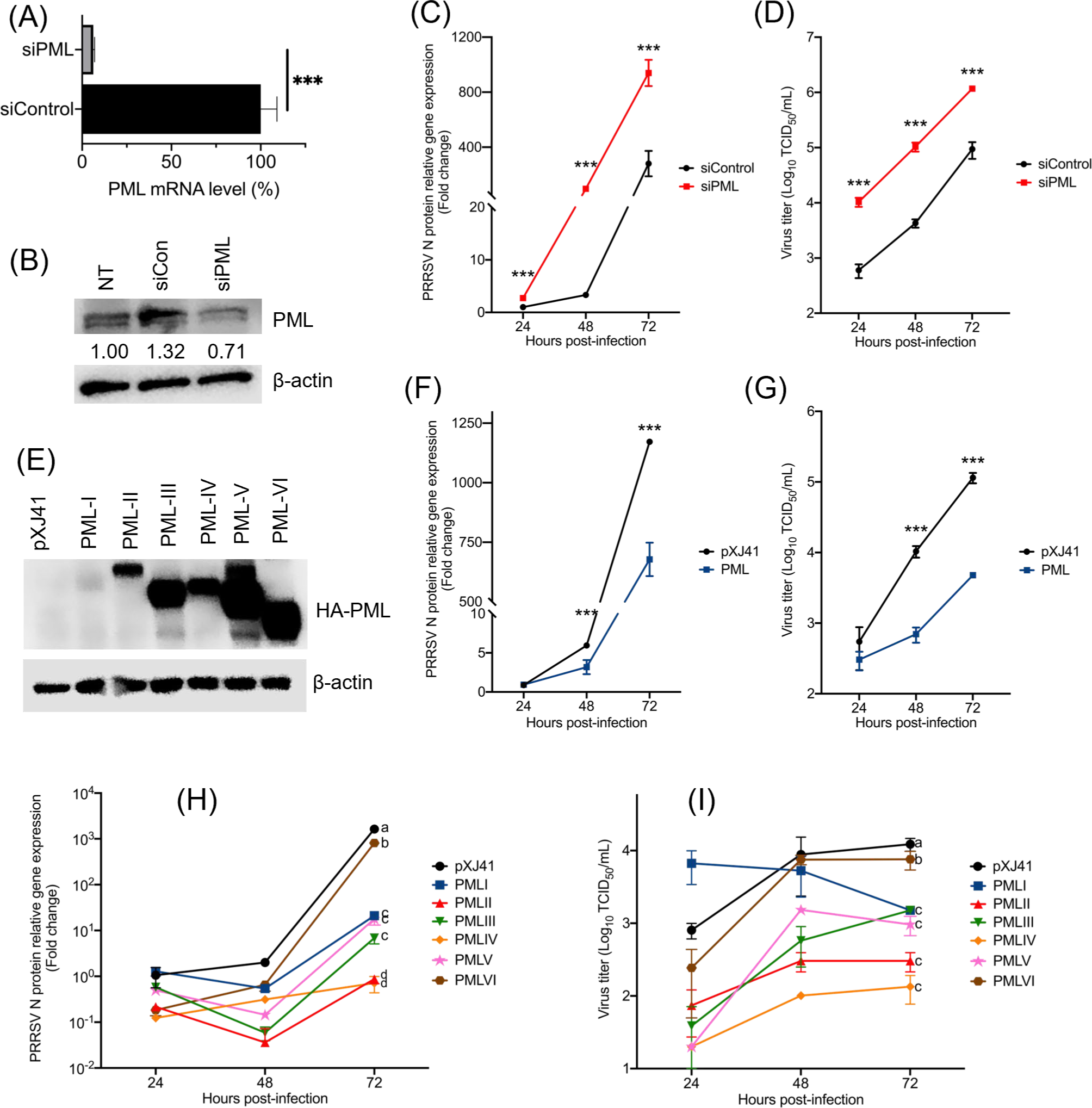
Restriction of PRRSV replication by PML protein. siRNA-mediated gene silencing of PML. MARC-145 cells were transfected with siControl or siPML, and relative amounts of protein were determined at 48 h post-transfection. (A), PML mRNA were measured by RT-qPCR using specific primers. (B), PML protein expression was analyzed by Western blot using α-PML PAb. β-actin served as a loading control. The intensities of PML were quantified with the Image J system, normalized to that of β-actin, and compared to the nontreated (NT) control. MARC-145 cells were treated with siRNA for 48 h and infected with PRRSV at an moi of 1. Supernatants were harvested at 0, 24, 48, and 72 hpi, and viral titers were determined by RT-qPCR (C) using primers specific for PRRSV N gene or 50% tissue culture infectious dose (TCID_50_) (D) as described in Materials and Methods. The error bars represent means ± standard deviation (s.d.) (n = 3). ***, P<0.001. (E), MARC-145 cells were expressed with individual HA-tagged PML isoforms and were analyzed by Western blot using α-HA MAb at 48 h post-transfection. Plasmids containing individual PML isoforms were equally mixed and expressed in MARC-145 cells for 48 h followed by infection with PRRSV at an moi of 1. Supernatants were harvested at indicated time, and viral titers were determined by RT-qPCR (F) or TCID_50_ assay (G). The error bars represent means ± standard deviation (s.d.) (n = 3). ***, P<0.001. (H, I) MARC-145 cells were expressed with individual PML isoforms for 48 h, followed by infection with PRRSV at an moi of 1. Supernatants were harvested and viral titers were determined by RT-qPCR (H) or TCID_50_ (I). The mean titers marked by lowercase letters show statistical significance (P < 0.05).

Since PML exhibited anti-PRRSV activity and because previous studies showed that different isoforms of PML had distinct biological roles (49), it was of interest to examine if a particular isoform is more potent than others for inhibition of PRRSV replication. To do this, each of the PML-I through PML-VI isoforms was over-expressed individually (Fig. 7E), and PRRSV titers were determined (Figs. 7H, 7I). While over-expression of PML-VI caused a slight reduction of N-specific mRNA, all other isoforms presented a significant decrease of N mRNA. Of these, PML-II and PML-IV caused the most drastic reduction at 72 hpi. Similar results were obtained for viral titers. PML-VI caused a moderate level of reduction in the viral titer compared to control. In contrast, the viral titers were reduced by 1.52 logs and 1.88 logs in cells over-expressing PML-II and PML-IV, respectively, at 72 hpi. Our results indicate that PML exhibited the antiviral function and inhibited PRRSV replication.

### PML interactions with nsp1 of other arteriviruses

PRRSV is a member of the family *Arteriviridae*, and thus, it was of interest to examine whether the PML downregulation by nsp1 protein was a common strategy for immune evasion for other arteriviruses. Besides PRRSV, equine arteritis virus (EAV), lactate dehydrogenase elevating virus (LDV) of mice, and simian hemorrhagic fever virus (SHFV) are the most studied member viruses in the family, and thus nsp1 protein of these viruses was examined. As with PRRSV nsp1, LDV nsp1 contains both PLP1α and PLP1β activities and thus is cleaved to two subunits nsp1α and nsp1β. While EAV nsp1 remains uncleaved due to the specific mutation for PLP1α, SHFV produces two subunits, nsp1αβ and nsp1γ. Thus, PRRSV nsp1β, LDV nsp1β, EAV nsp1, and SHFV nsp1αβ were individually expressed and examined for the suppression of PML-NB formation in the nucleus (Fig. 8A, 8B). All of PRRSV nsp1β, LDV nsp1β, EAV nsp1, and SHFV nsp1αβ decreased the number of PML-NBs in the nucleus compared to control after IFN stimulation. Co-localization of SHFV nsp1αβ and PML-NBs was observed in the nuclei of SHFV nsp1αβ-expressing cells (Fig. 8A), indicating the share of the binding between PML and nsp1 of other arteriviruses. The co-IP assay demonstrated that PML co-precipitated with PRRSV nsp1β, LDV nsp1β, EAV nsp1 as well as SHFV nsp1αβ (Fig. 8C). PML, however, did not precipitate LDV nsp1α and PRRSV nsp1α, which is consistent with the previous finding in Fig. 5B. Taken together, this study shows that the PML degradation by nsp1 is a common strategy of arteriviruses for the immune evasion and enhancement of viral replication.

**Fig. 8.**
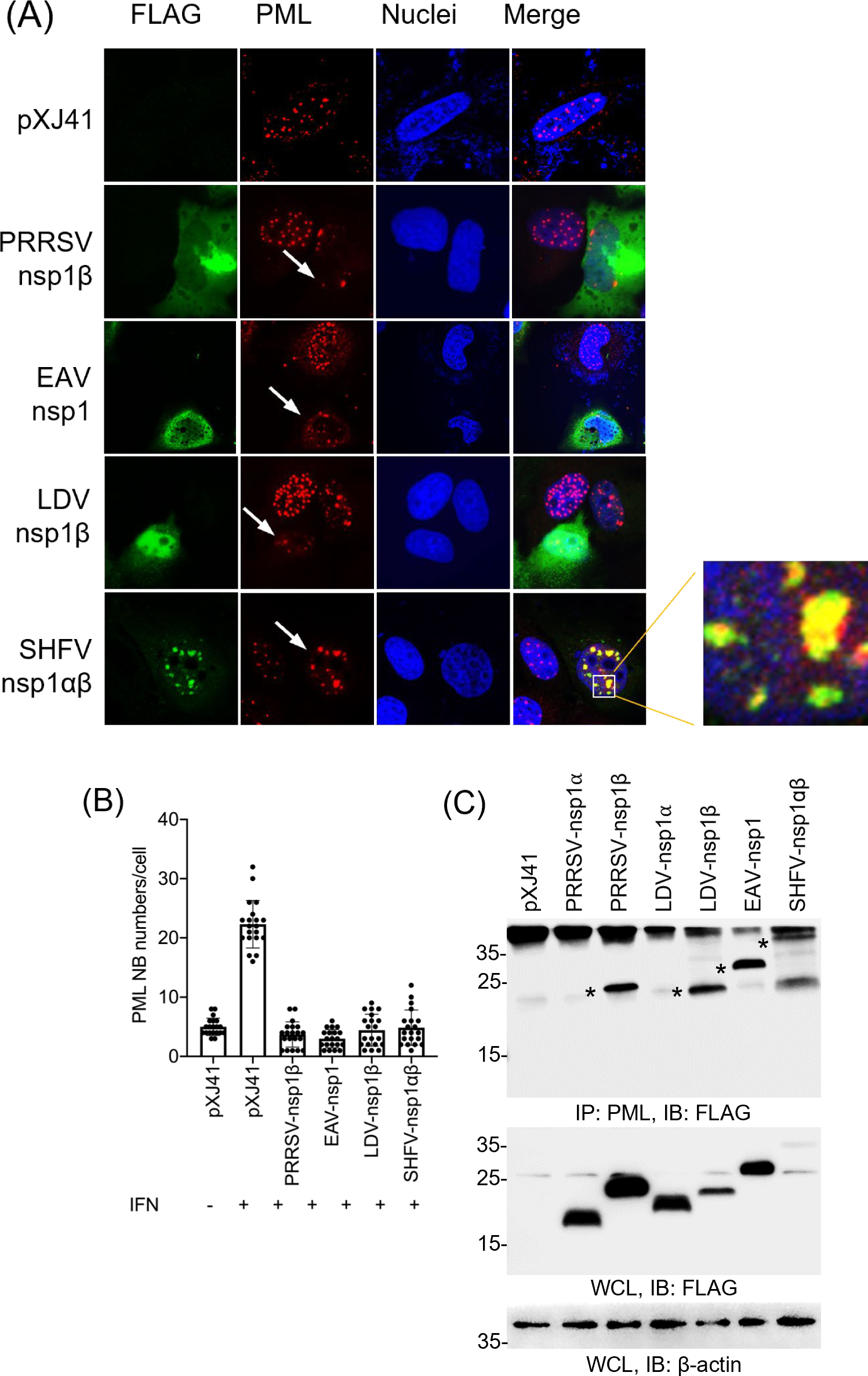
PML protein binding by nsp1 subunits of different arteriviruses. (A), HeLa cells were transfected with indicated FLAG-tag arterivirus nsp1 subunit genes and stimulated with IFN-β (1000 unit/ml). The cells were stained with α-FLAG MAb (green) and α-PML PAb (red). Nuclei were stained with DAPI (blue). The white box indicates the colocalization of nsp1β and PML, and the partially enlarged image is demonstrated. The images were taken by confocal microscopy (Nikon A1R). (B), The PML-NB numbers in each indicated gene-transfected cell nuclei. The PML-NB puncta were counted in randomly chosen 20 nuclei. (C), Specific binding of nsp1β to endogenous PML determined by co-IP. HeLa cells were co-transfected with 2 μg of indicated genes for 24 h and were subjected to co-IP. Cells lysates were pulled down by α-PML PAb and were probed with α-FLAG PAb. Whole cell lysates (WCL) were subjected to Western blot as an input control. Stars indicated specific proteins co-precipitated by PML

## DISCUSSION

Promyelocytic leukemia (PML) protein is a member protein of the TRIM family. It was initially identified as a hybrid protein containing retinoic acid receptor α which was associated with acute promyelocytic leukemia patients (50, 51). The TRIM protein superfamily, which PML belongs to, encompasses various restriction factors that serve as the first line of defense against invading pathogens. PML-NBs are stress-responsive nuclear structures and exhibit dynamic changes in size and composition depending on the cellular environment (22). These structures are crucial for cells to adapt to environmental cues and to maintain cellular homeostasis. Acting as a frontline defense, PML protein responds to viral infections. Conversely, viruses have evolved to counter-measure the host antiviral defense and to modify PML-NBs, creating a cellular environment conductive to viral replication. In the present study, we have uncovered the underlying mechanism for interactions between the swine arterivirus PRRSV and host cell protein PML and established a working model (Fig. 9).

**Fig. 9.**
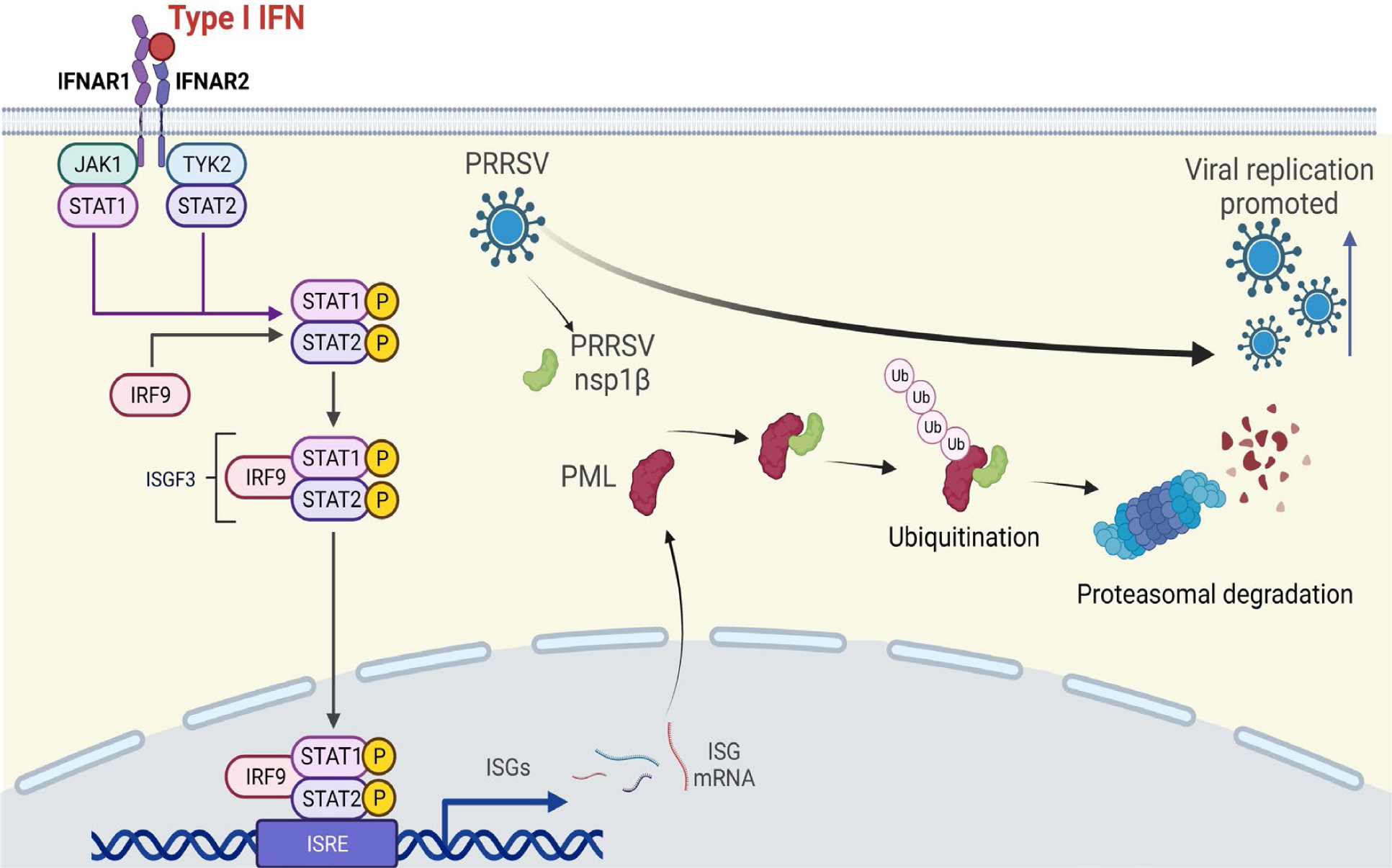
Working model for nsp1β-mediated PML degradation and promotion of viral replication. Virus infection typically triggers the immediate production of type I interferons (IFNs-α/β) in infected-cells. IFNs are secreted out from infected cell and bind to IFN receptors on the surface of neighbor cells. JAK and TYK kinases are activated and phosphorylate STATs which subsequently recruits IRF9 to form the ISGF3 complex. The complex is translocated to the nucleus, where binds to ISRE to express hundreds of antiviral proteins (ISGs) including PML. In PRRSV-infected cells, the nsp1β protein is translated from the incoming viral genome, and binds to PML. It is speculated that nsp1β may act as an E3 ligase for ubiquitination. Binding of nsp1β to PML leads to PML ubiquitination and proteasomal degradation. Downregulation of the antiviral PML protein is favorable for viral replication, which will eventually lead to the promotion of PRRSV replication. Images were created with BioRender.com.

Type I IFNs play an important role in the host innate immunity responding to viral infection, and PML is one of the ISGs expressed in response to infection. PML expression is IFN inducible, which is likely due to the presence of IFN-α-stimulated response elements (ISRE) and IFN-γ activation sites in the PML promoter (52). This indicates that PML acts as a downstream effector of IFNs. On the contrary, PRRSV nsp1β exhibits multifunctional properties to suppress type I IFN response during infection. Nsp1β can inhibit both the IFN production and IFN signaling pathways and results in the suppression of ISG expressions including PML protein. Although PML is a nuclear protein and many RNA viruses replicate in the cytoplasm, multiple studies show that even RNA viruses have mechanisms to counteract the expression and function of PML protein (40, 41, 44, 53–56). PML downregulation, however, does not exclude viral inhibition through the upstream IFN signaling pathway which leads to the suppression of IFN and ISG expressions(41). Previously, we have shown that nsp1β can block the host mRNA nuclear export and downregulates host gene expressions (19, 20). Nevertheless, the PML mRNA transcription was not decreased in nsp1β-expressing cells (Fig. 2). This finding suggests that PML downregulation is likely post-translational than transcriptional. Interestingly, we find that nsp1α also decreases the PML-NB formation and downregulates the PML expression. PRRSV nsp1α has been shown to downregulate type I IFN response through different mechanisms from nsp1β (12, 14–16, 57, 58). Our study presents initial findings that nsp1α directly contributes to ISG modulation. Moreover, the PML downregulation by two viral proteins suggests a pivotal role of PML in the antiviral response. The virus has seemingly evolved to equip diverse mechanisms to counteract the antiviral function exerted by PML, highlighting its significance in host-virus interactions.

Previous studies provide insights into various mechanisms employed by RNA viruses to degrade PML protein. One such mechanism involves the degradation of PML induced by viral proteases as shown for encephalomyocarditis virus and enterovirus 71 (40, 41). For these viruses, 3C^pro^ cleaves the isoforms of PML or alters the cellular distribution of PML and reduces its expression. In the case of PRRSV, the protease activity of nsp1β was initially thought to involve in PML degradation. The mutational studies, however, show that PLP1β is not involved in the decrease of PML-NB formation. A previous study shows that once nsp1β is cleaved from nsp2, the PLP1β domain in nsp1β undergoes conformational changes and becomes inactive due to the self-blocking in its C-terminal extension (59), supporting our finding. HSV-1, HTLV-1, and EMCV infections also trigger degradation of PML, and this degradation process relies on the activity of the proteasome (32, 40, 55, 56, 60). In our study, nsp1β protein governs the regulation of PML function through ubiquitin-dependent proteasomal degradation. From the colocalization studies, nsp1β was postulated to interact with PML protein, and this interaction was confirmed by the GST pulldown and co-IP assays. Further investigation demonstrates two SIM motif of nsp1β as the PML-binding domain. The SIMs and nsp1β interactions facilitated by SUMOylation of PML plays a pivotal role in modulating various functions, including the formation of macromolecular structures such as PML-NBs (61). Our study shows that the interaction between SUMO of the PML protein and the SUMO-interacting motif of nsp1β is crucial for PML degradation (Fig. 6). PML is a critical component in initiating the IFN response by facilitating the assembly of transcription complexes on the IFN-β promoter (33), and our study demonstrates that the PML degradation by PRRSV nsp1β results in the suppression of IFN-β production. This effect is dependent on the IRF3 activity (Fig. 6E). Additionally, the nsp1 subunit of other arteriviruses also contains a similar function of PML downregulation (Fig. 8). The binding of LDV nsp1β, EAV nsp1, and SHFV nsp1αβ to PML established a common mechanism for PML regulation by different arteriviruses.

The PML protein has been implicated to mediate antiviral responses against a wide range of RNA viruses through diverse mechanisms (34, 36, 41, 60, 62–64). In our study, PML gene-silencing enhanced PRRSV replication, indicating that the PML downregulation is advantageous for PRRSV replication. PML-II specifically interacts with the transcription factors NF-κB, STAT1, and CBP, and facilitates the formation of transcription complexes on the promoters of IFN-β and numerous ISGs, thereby enabling efficient transcription (65). PML-IV has been identified as an inhibitor of vesicular stomatitis virus through the augmentation of IRF3 activation (34). In our study, overexpression of PML-II and PML-IV also caused the most significant inhibition of PRRSV replication, further supporting the critical role of PML-II and PML-IV in exerting antiviral activities.

In summary, our research has revealed that the PML protein plays a key role in restricting the replication of PRRSV. To counteract this antiviral function, viral nsp1β binds to PML and mediates its degradation. Our study sheds light on how nsp1β of the swine arterivirus PRRSV promotes its benefit by degrading PML.

## MATERIALS AND METHODS

### Cells and viruses

HeLa (NIH HIV Reagents Program, Germantown, MD) and MARC-145 cells were maintained in Dulbecco’s modified Eagle’s medium (DMEM; Corning Inc., Corning, NY) supplemented with 10% heat-inactivated fetal bovine serum (FBS; Gibco, Grand Island, NY), in a humidified incubator with 5% CO2 at 37 °C. Anti (α)-FLAG PAb (rabbit) was purchased from Rockland Inc (Gilbertsville, PA). α-p65 Mab (mouse) (F-6, sc-8008) was purchased from Santa Cruz Biotechnologies Inc. (Santa Cruz, CA). Alexa-Flour 488-conjugated, and Alexa-Flour 568-conjugated secondary antibodies were obtained from ThermoFisher (Rockford, IL). Human tumor necrosis factor-α (TNF-α) (8902) was purchased from Cell Signaling (Danvers, MA). DAPI (4′, 6′-diamidino-2-phenylindol) was obtained from Sigma (St. Louis, MO). North American genotype PRRSV strain PA8 was propagated in MARC-145 cells and used for this study. For infection, cells were infected at a multiplicity of infection (moi) of 1. MARC-145 cells were grown to approximately 70% confluence for gene transfection or virus infection. Viral titers were determined by a 50% tissue culture infectious dose (TCID_50_) assays in MARC-145 cells.

### Antibodies and chemicals

Antibodies and chemicals used in the present study are listed as follow. Anti (α)-PRRSV-N protein MAb (MR40; mouse monoclonal antibody) was obtained from E. Nelson (South Dakota State University, Brookings, SD). α-PML PAb (rabbit) (sc-5621), α-β-actin MAb (mouse) (C4; sc-47778), and α-Ubi MAb (mouse) (P4D1: sc-8017) were purchased from Santa Cruz Biotechnologies Inc. (Santa Cruz, CA). α-FLAG PAb (rabbit) was purchased from Rockland Inc. (Gilbertsville, PA). α-FLAG MAb (rat) (M2) was purchased from Agilent (Santa Clara, CA). α-HA MAb (mouse), Alexa-Fluor 488-conjugated, and Alexa-Fluor 568-conjugated secondary antibodies were purchased from ThermoFisher (Rockford, IL). Polyinosinic:polycytidylic [poly (I:C)], and DAPI (4=,6-diamidino-2-phenylindole) were purchased from Sigma (St. Louis, MO). MG132 was purchased from Calbiochem (San Diego, CA). Human interferon-β recombinant protein was purchased from Millipore (St. Louis, MO), and for stimulation, 1000 unit/ml was added to cells for 6 h.

### Genes and plasmids

The genes for nsp1β and N with FLAG tag were cloned from PRRSV-2 strain VR2332 and inserted into the pXJ41 expression plasmid as described previously (20). FLAG-nsp1α and FLAG-nsp1β of LDV, FLAG-nsp1 of EAV, nsp1αβ of SHFV, and ORF3a of SARS-CoV-2 were cloned into pXJ41-expressing vectors as described previously (20, 38, 39). The mutant plasmids nsp1β-K124A, nsp1β-L126A, nsp1β-C90A, and nsp1β-H159A were constructed by PCR-based site-directed mutagenesis as described elsewhere (4). The HA-PML-I, HA-PML-II, HA-PML-III, HA-PML-IV, HA-PML-V, and HAPML-VI genes were subcloned from the plasmid phNGX-PMLI-VI into the pXJ41 expression vector using the Hind III and Xho I recognition sequences. The pcDNA-HA-Ubiquitin was provided by Ying Fang (University of Illinois at Urbana-Champaign, Urbana, IL). The pIFN-β-luciferase reporter plasmid and pIRF3-luciferase reporter plasmid were provided by Stephan Ludwig (Institute of Molecular Medicine, Heinrich Heine Universtät, Düsseldorf, Germany). The pRL-TK Renilla luciferase reporter plasmid was purchased from Promega (Madison, WI). E. coli strain DH5α was used for transformation.

### DNA transfection and dual luciferase assay

DNA transfection was performed using Lipofectamine 200 according to the manufacturer’s instruction (Invitrogen). Cells were grown in 12-well plates and transfected with 0.5 μg of viral gene, 0.5 μg of reporter plasmid, and 0.05 μg of pRL-TK in 1:1:0.1 ratio. 0.5 μg of poly(I:C) was transfected to cells for stimulation for 16 h at 24 h after DNA transfection. Cell lysates were prepared using Passive lysis buffer (Promega). Supernatants were collected and measured for luciferase activities using the Dual luciferase reporter assay system (Promega). Signals were determined in the luminometer (Wallac 1420 VICTOR multi-label counter, Perkin Elmer, Waltham, MA). Values for firefly luciferase reporter activities were normalized by the Renilla internal control, and results were expressed as relative luciferase activities. The assay was repeated twice, and each assay was conducted in triplicate.

### Immunofluorescence analysis (IFA)

HeLa cells were grown on microscope coverslips and fixed with 4% paraformaldehyde in phosphate-buffered saline (PBS) for 1 h at room temperature (RT), followed by three washes with PBS. The cells were permeabilized with 0.1% Triton X-100 for 15 min at RT, followed by three washes with PBS. After incubation with 1% bovine serum albumin (BSA) in PBS for 1 h at RT, cells were incubated with a primary antibody (1:200) in blocking buffer for 2 h, followed by three washes with PBS and incubation with a secondary antibody (1:200) for 1 h. The cells were stained with DAPI (1:5,000) for 5 min, and after a final wash with PBS, the coverslips were mounted on microscope slides using Fluoromount-G mounting medium (Southern Biotech, Birmingham, AL). The cells were examined using a Nikon A1R confocal microscope. To quantify reduction of PML, the formula described previously were followed (58).

### Reverse transcription-quantitative PCR (RT-qPCR)

Total cellular RNA was extracted using the TRIzol reagent according to the manufacturer’s instruction (Invitrogen). RT-qPCR was performed in the ABI sequence Detector System (ABI Prism 7000 Sequence Detection System and software: Applied Biosystems) using a final volume of 25 μl containing 2 μl of cDNA from reverse-transcription reaction, a primer mix (2.5 pM each of sense and antisense primers), 12.5 μl of SYBR Green Master Mix (Applied Biosystems), and 8 μl of distilled water. The primer sequences were listed as follow; for PML, forward 5’-CATCACCCAGGGGAAAGATG-3’, reverse 5’-GGTCAACGTCAATAGGGTCC-3’; for PRRSV-N, forward 5’-GATAACCACGCATTTGTCGTC-3’, reverse 5’-TTGAACAAATTAAAACAAAAAGGTG-3’, for GAPDH, forward 5’-CGGAGTCAACGGATTTGGTCGTA-3’, reverse 5’-AGCCTTCTCCATGGTGGTGAAGAC-3’. The amplification parameters were 40 cycles of two steps each cycle comprised of heating to 95 °C and 60 °C. The mRNA levels were calculated using the 2^−ΔΔCT^ method (66) and normalized using GAPDH.

### GST pulldown assay

pGEX-4T3-nsp1β was expressed in E. coli strain BL21 (DE3) as described previously (67). Briefly, bacteria were transformed with the plasmid and grown to an optical density at 600 nm of 0.6 and stimulated with IPTG (1 mM) for 3 h. The bacteria were collected, resuspended in cold PBS, and sonicated on ice three times for 30 s each time with 2 s intervals (Soniprep 150; Gallenkamp). The proteins were solubilized with 1% Triton X-100, and the supernatants were incubated with glutathione-Sepharose 4B beads (GE Healthcare). The beads were collected, washed with PBS, and finally resuspended in binding buffer (20 mM Tris-HCl, 100 mM KCl, 2 mM CaCl, 2 mM MgCl2, 5 mM dithiothreitol [DTT], 0.5% NP-40, 1 mM phenylmethylsulfonyl fluoride [PMSF], 5% glycerol). For GST pulldown assays, whole-cell lysates were prepared and incubated with GST fusion proteins in binding buffer at 4°C overnight. The beads were washed three times with binding buffer and boiled at 95°C for 10 min in SDS-PAGE sample buffer. Supernatants were collected by centrifugation at full speed in a microcentrifuge at 4°C for 5 min, and proteins were resolved by SDS-10% PAGE. The gels were subjected to Western blot assays.

### Co-immunoprecipitation (co-IP) and Western blot

Cell lysates were prepared in lysis buffer (50 mM Tris-HCl, 0.1% Triton-X-100, 150 mM NaCl, pH 7.8) containing 1X protease inhibitors (Thermo, Rockford, IL) by rotating at 4 °C for 30 min. Cell lysates were aliquoted to 100 μl as the whole cell lysates and stored at −80 C until use. For immunoprecipitation, cell lysates were incubated with antibody overnight at 4°C with gentle agitation. 30 μl of protein G agarose beads (EMD Millipore, Temecula, CA) was added, and the mixture was further incubated for at least 2 h at 4°C. The beads were washed three times with lysis buffer, and the precipitates were eluted in 30 μl of M-PER mammalian protein extraction reagent (Thermo) and 6x loading buffer by boiling at 95°C for 5 min. The beads were spun down in a microcentrifuge at full speed at 4°C for 5 min, and the supernatants were subjected to SDS-10% PAGE. The resolved proteins were transferred to an Immobilon-P polyvinylidene difluoride (PVDF) membrane (Millipore), and the membranes were incubated with Tris-buffered saline with Tween 20 (TBST) blocking buffer (10 mM Tris-HCl, 150 mM NaCl, 0.05% Tween 20 containing 5% skim milk powder) for 1 h at RT, followed by further incubation with primary antibody at 4°C overnight. The membranes were washed five times with TBST and incubated with peroxidase-conjugated secondary antibody in TBST blocking buffer for 1 h. The membranes were washed five times, and proteins were visualized using the ECL detection system (Thermo).

### siRNA-mediated PML knockdown

To knock-down the PML gene in MARC-145 cells, an siGenome human PML siRNA-Smart Pool (M-006547-01-0005) and an siGenome nontargeting siRNA pool (D-001206-13-05) were purchased from Dharmacon (Lafayette, CO). An siGenome nontargeting siRNA pool was used as a control with no specific gene targeting. MARC-145 cells were grown to 50% confluence in 6-well plates, and 100 pmol of siRNA per well was transfected using Lipofectamine 2000 (Invitrogen, Carlsbad, CA) according to the manufacturer’s instructions. Cell lysates were collected at 24 h or 48 h post-transfection and analyzed by RT-qPCR and Western blot to determine the knockdown efficiency.

### Statistical analysis

Statistical significance was determined by two-tailed Student’s t-test. Data analyses were performed using GraphPad Prism version 9.00 (San Diego California USA).

## Data availability

The authors confirm that the data supporting the findings of this study are openly available within the article. All other raw data will be shared upon request.

## ACKNOWLEDGMENTS

This project was supported by Agriculture and Food Research Initiative (AFRI) Competitive Grants nos. 2018-67015-28287 and 2023-67015-39710 from the U.S. Department of Agriculture (USDA) National Institute of Food and Agriculture (NIFA) awarded to DY.

